# Revamped transposon-induced hepatocellular carcinoma model reveals deep similarities of cancer-driver landscapes across mice and humans

**DOI:** 10.64898/2026.10.31.763279

**Authors:** Gergely Imre, Andrea Mátés-Nagy, May Raya, Olivia Orsolya Verebi, Liza Hudoba, Anna Faragó, Dóra Latinovics, Réka Karkas, Péter Germán, Péter Kálmán, Ede Migh, Aladár Pettkó-Szandtner, Orsolya Kolacsek, András Füredi, Péter Horváth, Tamás Orbán, István Nagy, Lajos Mátés

## Abstract

Genome-scale cancer-driver discovery is limited by statistical uncertainty in driver calling and by experimental constraints on mutations available for selection. We developed 3×fOnc3—a three-frame Sleeping Beauty (SB) mutagenesis system designed to broaden functional perturbations and reduce integration biases of traditional SB screens—and tested it in vivo in mouse liver. Within 12 months, 60% of mice developed heterogeneous hepatocellular tumors. Tumor-selected integrations were interpreted against an empirical reference from SB-based therapeutic gene delivery to hepatocytes, enabling correction for gene targetability. Background-aware filtering prioritized 315 driver candidates, including 243 protein-coding genes. Tumors were multiclonal, and identical integration-marked lineages expanded differently across lesions, supporting context-dependent clonal expansion. SB hits within young L1 subfamily members were selectively enriched in tumors. Human orthologues of candidate drivers were enriched for Cancer Gene Census genes and protein-altering pan-cancer mutations. Together, 3×fOnc3 reveals a background-corrected driver landscape linking driver selection to clonal tumor evolution and cross-species cancer relevance.

## Introduction

Cancer genome sequencing has revealed a vast landscape of somatic alterations, but distinguishing mutations that actively contribute to tumor development from the much larger passenger background remains a central challenge in cancer genetics. Computational driver-discovery methods address this problem by identifying mutational patterns that deviate from expectations under neutral mutagenesis and have generated increasingly comprehensive catalogues of positively selected cancer genes[1].

Nevertheless, statistical inference is influenced by gene-specific mutational opportunity, heterogeneous background mutation rates, cohort composition and the limited power to detect infrequent or context-dependent driver events. Hepatocellular carcinoma (HCC) exemplifies this complexity: genomic studies have identified alterations across multiple signalling and regulatory pathways, with substantial heterogeneity between tumors and combinations of recurrent and rare candidate drivers [2]. Indeed, even recent deep whole-genome analysis of nearly 500 HCCs identified additional coding and non-coding candidate drivers, indicating that the functional driver landscape remains incompletely resolved [3]. Experimental approaches that interrogate driver activity through in vivo selection can therefore provide information complementary to mutation-based inference from human tumor cohorts.

Insertional mutagenesis provides such a forward-genetic strategy because the evolving tumor itself selects genetic perturbations that confer a fitness advantage in the tissue in which they arise. The Sleeping Beauty (SB) transposon system established somatic insertional mutagenesis as a scalable approach for cancer-gene discovery in solid tissues [4, 5]. Mutagenic T2/Onc-family transposons combine bidirectional splice-acceptor/polyadenylation modules capable of disrupting genes with promoter/splice-donor elements capable of activating downstream gene sequences, thereby enabling both loss- and gain-of-function selection. Liver-restricted SB mutagenesis subsequently demonstrated the applicability of this approach to hepatocarcinogenesis: analysis of 8,060 nonredundant integrations from 68 tumor nodules identified 19 HCC-associated common insertion sites, including established cancer genes such as MET and EGFR [6]. These studies established the power of transposon-based forward genetics, but also highlighted the importance of experimental architecture in determining which genomic perturbations can be generated, selected and ultimately recovered.

Several features can limit the interpretation and functional reach of conventional insertional screens. First, recurrence is not determined exclusively by biological selection. In traditional SB cancer screens, mobilization from chromosomally integrated transposon concatemers additionally creates local hopping, a preferential reintegration of transposons into regions linked to the donor locus that can distort the genomic space sampled by mutagenesis. By contrast, SB launched from episomal vectors displays a substantially more dispersed genome-wide integration profile. Even in this setting, however, integration probabilities are not completely uniform and are influenced by target-sequence availability and local genomic and chromatin features [7]. Such intrinsic targeting biases can generate recurrent integration hotspots in the absence of tumor selection [8].

Consequently, highly targetable genes can accumulate integrations independently of tumor-promoting activity, making an experimentally matched background integration landscape particularly valuable for gene-level driver inference. Second, the mutational consequences accessible to selection depend on transposon design. In T2/Onc-type vectors, promoter-driven expression of truncated host proteins requires not only appropriate integration position and orientation but also productive translation of the downstream coding sequence. Thus, translational frame represents an additional constraint on the gain-of-function mutational space sampled by the screen. Finally, recurrence identifies loci repeatedly favoured during tumor development but does not by itself establish that a given mutation produces a fixed degree of clonal expansion. In genetically heterogeneous and multiclonal tumors, the realized representation of an integration-bearing lineage may also depend on additional alterations, interacting cellular populations and local tissue context. Separating gene-specific selection from integration-site accessibility, functional configuration and subsequent clonal behaviour therefore remains a major challenge for forward-genetic cancer screens.

Here, we developed 3×fOnc3, a somatically delivered three-frame SB insertional-mutagenesis system designed to broaden access to promoter-driven truncated-protein configurations while restricting transposition to an early period of tumor development. To distinguish tumor selection from intrinsic SB targetability, we compared the resulting integration landscape with a large, biologically matched reference generated by therapeutic SB transposition during Fah-driven hepatocyte repopulation. We combined background-adjusted gene recurrence with selection of truncated-protein-overexpression-compatible events to prioritize candidate cancer drivers, and used integration-site readouts to examine how recurrent driver-associated lineages were represented within and between spatially distinct tumors. Finally, we mapped the resulting protein-coding candidate set to human orthologues and assessed its cancer relevance using established cancer-gene annotations, pan-cancer somatic mutation data and human pathway information. Together, this framework links experimentally controlled forward genetics with clonal tumor evolution and cross-species cancer genomics to interrogate both the identity and the behaviour of somatically selected cancer-driver candidates.

## Results

### Creation and testing of the 3×fOnc3 somatic mutagenesis system

The T2/Onc3 transposon provides a versatile mutagenic architecture in which bidirectional splice-acceptor/polyadenylation modules can disrupt endogenous gene expression, whereas a strong promoter coupled to a splice donor can drive aberrant downstream transcripts [9]. To broaden the range of positively selectable mutations accessible to this system, we developed the 3×fOnc3 strategy around three principal features: somatic delivery of mutagenic transposons directly into hepatocytes, efficient chromosomal integration mediated by the hyperactive SB100 transposase, and three related T2/Onc3 constructs providing alternative translational reading frames (**Fig. 1A**). The three reading-frame variants were designed to minimize and standardize the vector-derived N-terminal sequence appended to truncated proteins, while extending promoter-driven truncated-protein configurations across all three reading frames. Translation from all three constructs begins with Met-Gly (**Fig. 1A**), and removal of the initiator methionine is expected to expose glycine as the mature N-terminal residue (See more details in Supplementary material 1). Each construct also carried a distinct two-nucleotide barcode (Supplementary material 1) within the left SB inverted terminal repeat (ITR), enabling direct identification of the transposed frame variant by the SBE-seq protocol [7] suitable for identifying transposon-genome junction sequences. Introduction of these barcodes did not detectably impair transposition in a cell-based assay (Supplementary material 1: **Fig. S1**).

**Figure 1.**
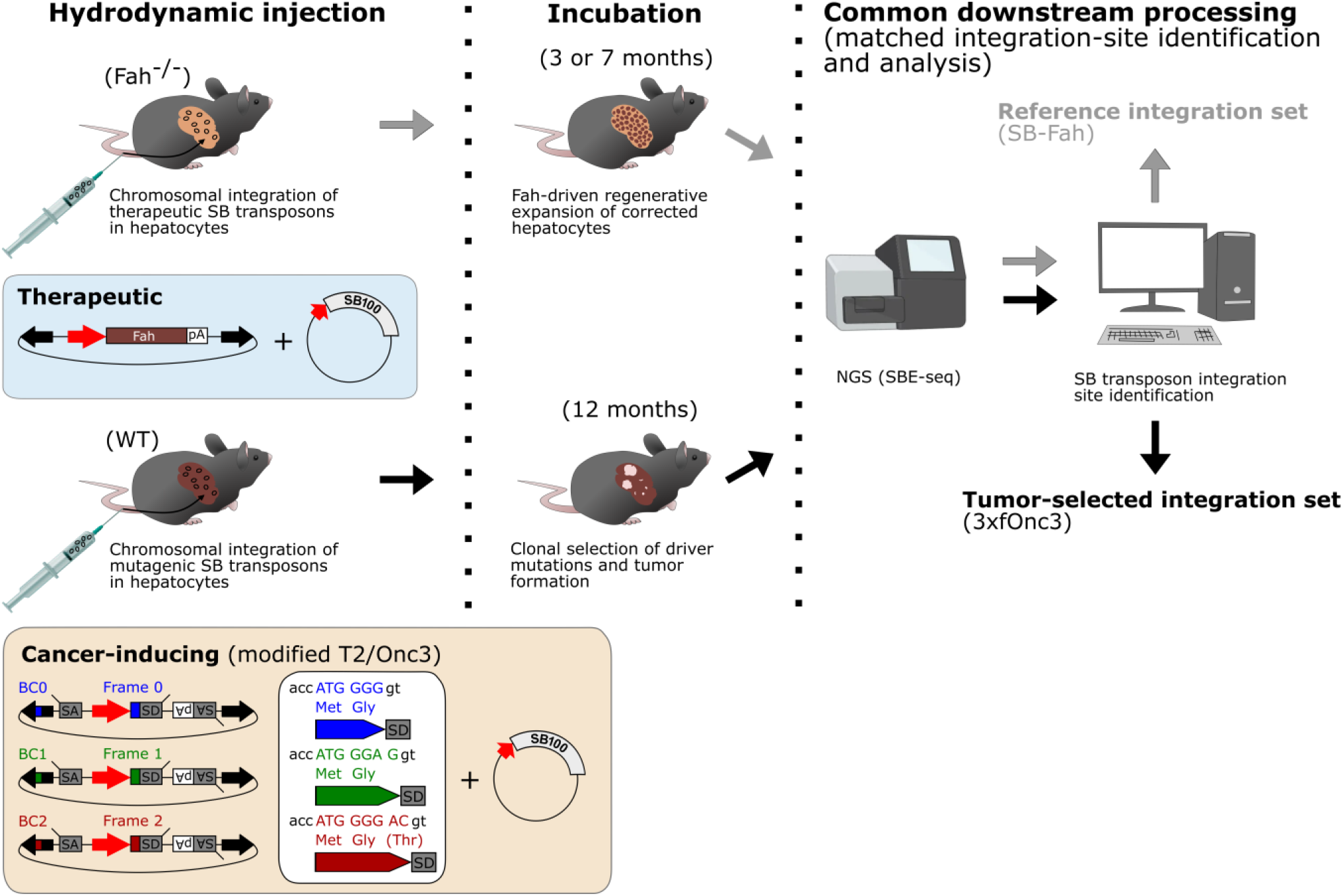
Design of the 3×fOnc3 somatic mutagenesis system and induction of heterogeneous hepatocellular tumors in mice. Schematic overview of the experimental framework. In the reference arm (top), Fah-deficient (Fah^−/−^) mice received hydrodynamic delivery of a therapeutic SB transposon plasmid carrying a *Fah* therapeutic minigene together with a plasmid expressing the hyperactive SB100 transposase. At 3 or 7 months post injection, following Fah-driven regenerative expansion in the absence of tumorigenic selection, whole-liver samples were used to generate the SB–Fah reference integration set. In the tumor-induction arm (bottom), wild-type mice received a mixture of three modified T2/Onc3 transposon plasmids together with the same SB100 transposase-expression plasmid. The three 3×fOnc3 variants provide alternative translational reading frames (frames 0, 1 and 2) and carry distinguishable two-nucleotide barcodes within the left SB inverted terminal repeat; the frame-0 construct retains the wild-type left-ITR sequence. Following approximately 12 months of in vivo selection and tumor development, 3×fOnc3 samples were processed to generate the tumor-selected integration set. SB–genome junctions from both experimental arms were analysed by SBE-seq followed by matched integration-site identification and downstream computational analysis. SA, splice acceptor; SD, splice donor; pA, polyadenylation signal.

To provide a biologically matched reference for subsequent integration-site analyses, we incorporated a previously established therapeutic SB-mediated fumarylacetoacetate hydrolase (Fah) gene transfer dataset into the experimental framework (**Fig. 1A**). In Fah-deficient mice, hydrodynamic delivery of a therapeutic Fah transposon together with the same hyperactive SB100 transposase resulted in chromosomal integration followed by Fah-driven regenerative expansion of corrected hepatocytes in the absence of tumorigenic selection [7]. The SB–Fah material therefore provided a closely matched non-tumorigenic reference for comparison with the 3×fOnc3 tumor-induction system.

Hydrodynamic delivery was used to introduce the 3×fOnc3 constructs into 30 two-month-old C57BL/6J mice (15 males and 15 females). We mixed the F0, F1, and F2 3×fOnc3 plasmids at 45%, 30%, and 25%, respectively, to approximate the unequal representation of the corresponding intron phases in the mouse genome [10]. Previous analysis of the corresponding somatic SB delivery system showed that detectable transposition ceases within approximately one week, restricting mutagenesis predominantly to an early burst [7]. After approximately 12 months of in vivo selection, macroscopically detectable liver tumors developed in 18 of 30 animals (60%), including 12 of 15 males (80%) and 6 of 15 females (40%). The resulting liver tumors displayed substantial macroscopic heterogeneity in tumor burden, size and morphology across individual animals (Supplementary material 1: **Fig. S2A**). Histopathological examination confirmed hepatocellular tumor morphology, while immunohistochemical staining for hepatocellular and HCC-associated markers further supported the hepatocellular identity of the lesions (Supplementary material 1: **Fig. S2B**). Thus, an early burst of 3×fOnc3 mutagenesis was sufficient to generate a heterogeneous spectrum of hepatocellular tumors during prolonged in vivo selection.

### Somatic 3×fOnc3 mutagenesis reveals distinct selection of protein-coding and L1-associated integration landscapes during liver tumorigenesis

A central feature of the analysis was the large, biologically matched SB–Fah integration reservoir generated in mouse hepatocytes using the same SB100 transposase and somatic delivery route [7]. Because chromosomal integration was followed by Fah-driven regenerative expansion without tumorigenic selection, this dataset provides an empirical measure of the SB integration landscape generated in mouse hepatocytes under closely matched somatic delivery conditions. Unlike the theoretical genomic TA target space, it incorporates hepatocyte-specific genomic accessibility together with the intrinsic targeting preferences of the SB system. We therefore reprocessed the SB– Fah integration sites using the same computational framework applied to the 3×fOnc3 dataset (**Fig. 1A**).

To define the integration landscape underlying 3×fOnc3-induced liver tumorigenesis, we sequenced SB–genome junctions using SBE-seq. The final sequencing panel comprised 30 libraries from 17 animals: 16 individually dissected tumors, 10 pooled tumor samples and four whole-liver samples, of which two contained macroscopically visible tumors and two were macroscopically tumor-free (Supplementary Data D1). After integration-site calling and quality-control filtering, the final 3×fOnc3 dataset comprised 25,106 sample-level calls corresponding to 22,193 animal-level integration events; detailed filtering procedures are described in Methods.

The resulting integration landscape differed markedly from the SB–Fah reference. Individual Fah-corrected whole livers contained a median of 78,811 accepted SB integrations (n = 12), compared with 1,164.5 in macroscopically tumor-free 3×fOnc3 whole livers (n = 2), 1,033 in tumor-bearing whole livers (n = 2) and 534 in individually dissected tumors (n = 16) (**Fig. 2A**). This comparison indicates that only a small fraction of the initially generated integration landscape remained detectable after prolonged tumor selection, consistent with a much larger continuum of low-abundance integration-bearing lineages extending below the SBE-seq detection threshold.

**Figure 2.**
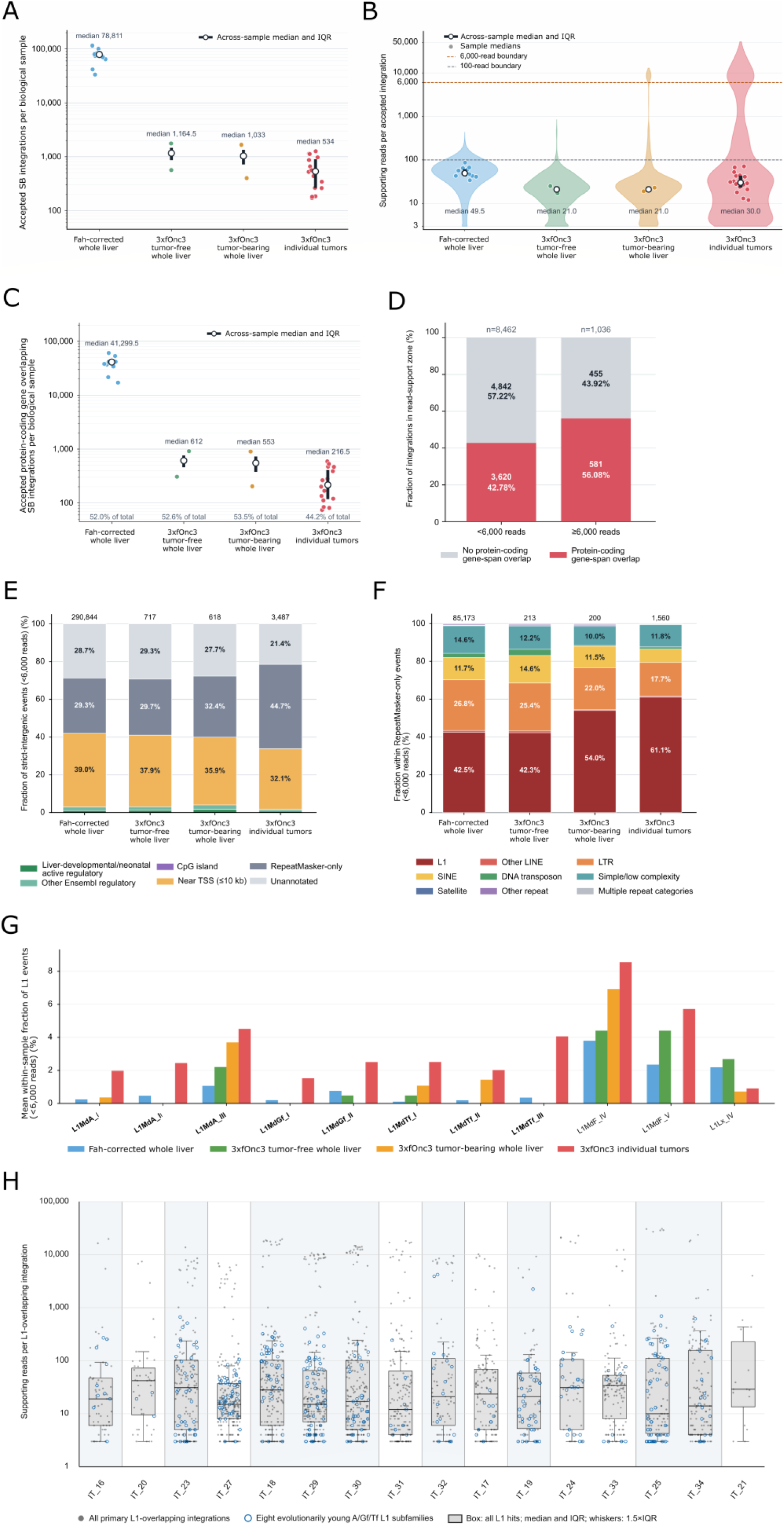
Tumor selection reshapes the 3×fOnc3 integration landscape and preferentially enriches low-support integrations in selected L1 subfamilies. (A) Number of accepted SB transposon integrations per biological sample in Fah-corrected whole livers (n = 12), macroscopically tumor-free 3×fOnc3 whole livers (n = 2), tumor-bearing 3×fOnc3 whole livers (n = 2), and individually dissected 3×fOnc3 tumors (n = 16). Colored points represent individual biological samples; open symbols and error bars indicate the across-sample median and interquartile range (IQR). Group medians are indicated above the distributions. The y-axis is logarithmic. (B) Sample-balanced distributions of supporting-read counts for accepted integration events in the same four sample groups. To prevent samples containing large numbers of integrations from dominating the group distributions, each biological sample contributed 2,000 deterministic quantiles and therefore equal weight to the corresponding violin. Gray points indicate the median read support of individual samples; open symbols and error bars indicate the across-sample median and IQR. Horizontal dashed lines mark the 100-read reference boundary and the empirically defined 6,000-read boundary separating the tumor-specific upper-support zone. Group median read supports are indicated below the distributions. The y-axis is logarithmic. (C) Number of accepted integrations overlapping protein-coding gene spans per biological sample. Symbols are as in panel A. The percentage shown below each group indicates the fraction of all accepted integrations in that group that overlapped a protein-coding gene. The y-axis is logarithmic. (D) Protein-coding gene overlap among integration events from the 16 individually dissected 3×fOnc3 tumors after stratification by read support. Bars show the fraction of integrations overlapping or not overlapping a protein-coding gene among events supported by <6,000 reads (n = 8,462) or ≥6,000 reads (n = 1,036). Counts and percentages are shown within each segment. Protein-coding gene overlap was enriched in the ≥6,000-read group (odds ratio = 1.71; two-sided Fisher’s exact P = 6.17 × 10⁻¹⁶). (E) Genomic annotation of strict-intergenic integrations supported by <6,000 reads. Events from each sample group were assigned to mutually exclusive annotation categories comprising liver developmental/neonatal active regulatory regions, other Ensembl regulatory regions, CpG islands, regions within 10 kb of a transcription start site (TSS), RepeatMasker-only annotations, and otherwise unannotated sequence. Numbers above bars indicate the total number of strict-intergenic events included in each group. Values represent fractions of pooled events within the indicated sample group. (F) RepeatMasker-class composition of the RepeatMasker-only events shown in panel E. Events were classified as L1, other LINE, LTR, SINE, DNA transposon, simple/low-complexity sequence, satellite, other repeat, or multiple repeat categories. Numbers above bars indicate the total number of RepeatMasker-only events in each group. Values represent fractions of pooled events within the indicated sample group. (G) Representation of evolutionarily young L1 subfamilies belonging to the currently active A, Gf and Tf lineages and selected older comparator subfamilies among L1-overlapping integrations supported by <6,000 reads. Bars show the sample-balanced mean within-sample fraction of primary L1-overlapping events assigned to each subfamily in Fah-corrected whole livers, macroscopically tumor-free 3×fOnc3 whole livers, tumor-bearing 3×fOnc3 whole livers, and individually dissected 3×fOnc3 tumors. The eight evolutionarily young A/Gf/Tf subfamilies are shown in bold on the x-axis. Differences between SB–Fah whole livers and individual 3×fOnc3 tumors were evaluated using 50,000 sample-label permutations; two-sided empirical P values were adjusted across tested subfamilies using the Benjamini–Hochberg procedure. FDR-adjusted q ≤ 0.05 was considered significant. (H) Supporting-read distributions of primary RepeatMasker L1-overlapping integrations in the 16 individually dissected 3×fOnc3 tumors. Gray points indicate all primary L1-overlapping integrations, whereas open blue circles indicate integrations assigned to one of the eight evolutionarily young A/Gf/Tf L1 subfamilies. Boxplots summarize the distribution of all L1-overlapping integrations within each tumor (median and IQR; whiskers, 1.5×IQR). Background blocks delineate tumor samples isolated from the same animal. Sample identifiers and corresponding animal and sequencing information are provided in Supplementary Data D1. The y-axis is logarithmic.

SB–Fah integrations showed a relatively compact read-support distribution, whereas 3×fOnc3 tumors contained an additional high-support component extending over several orders of magnitude (**Fig. 2B**). An empirical boundary of 6,000 reads defined an upper-support zone absent from the SB– Fah reference but present in all 16 individually analysed tumors. The median fraction of accepted integrations above this threshold was 11.68% in tumors compared with 0% in SB–Fah livers (two-sided Mann–Whitney P = 3.88 × 10⁻⁶). Additional distributional and RPM-normalized analyses independently confirmed this separation (Supplementary material 1). Supporting-read counts were therefore treated as a semiquantitative measure of clonal representation rather than converted directly into absolute cell numbers. The tumor-specific high-support component is thus most consistent with strongly expanded clones superimposed on a much broader population of lower-abundance integration-bearing lineages.

We next asked whether strong clonal representation was associated with preferential selection of integrations affecting protein-coding genes. At the whole-sample level, protein-coding gene-overlapping integrations accounted for 52.0% of events in SB–Fah whole livers, 52.6% in macroscopically tumor-free 3×fOnc3 whole livers and 53.5% in tumor-bearing whole livers, but only 44.2% in individually dissected tumors (**Fig. 2C**). Stratification by read support resolved this apparent paradox. Among 8,462 integrations supported by <6,000 reads, 3,620 (42.78%) overlapped a protein-coding gene, whereas 581 of 1,036 events (56.08%) in the ≥6,000-read zone did so (odds ratio, 1.71; two-sided Fisher’s exact P = 6.17 × 10⁻¹⁶; **Fig. 2D**). Thus, protein-coding gene overlap was specifically enriched among strongly expanded integration-bearing clones. This association does not imply that low-support protein-coding integrations are passengers, because read support reflects realized clonal representation rather than intrinsic driver potential alone.

To determine which genomic compartments accounted for the reduced protein-coding fraction within the low-support tumor population, we restricted subsequent analyses to strict-intergenic integrations supported by <6,000 reads. RepeatMasker-only annotations represented 44.7% of the 3,487 strict-intergenic low-support events in individual tumors, compared with 29.3% in the corresponding SB– Fah events (**Fig. 2E**). This shift was driven predominantly by LINE-1 (L1)-overlapping integrations: L1 elements accounted for 42.5% of RepeatMasker-only events in SB–Fah livers but 61.1% in individually dissected tumors (**Fig. 2F**), identifying endogenous L1 loci as a major component of the expanded low-support intergenic compartment.

We therefore examined whether this tumor-associated increase reflected L1 elements generally or particular evolutionary lineages. Eight evolutionarily young mouse L1 subfamilies belong to the currently active A, Gf and Tf lineages: L1MdA_I–III, L1MdGf_I–II and L1MdTf_I–III. Subfamily-level analysis revealed a highly non-uniform redistribution between the SB–Fah reference and 3×fOnc3 tumors (**Fig. 2G**). Six subfamilies were significantly enriched after correction for multiple testing—L1MdA_II, L1MdA_III, L1MdGf_II, L1MdTf_III, L1MdF_IV and L1MdF_V—whereas the evolutionarily older L1Lx_IV subfamily was significantly depleted. Four of the six enriched subfamilies therefore belonged to the evolutionarily young A, Gf or Tf lineages, with L1MdTf_III showing a particularly pronounced reservoir-to-tumor shift.

The association with young L1 lineages was confined to the low-support integration compartment. Among 3,487 strict-intergenic events supported by <6,000 reads, 1,268 (36.36%) overlapped an L1 element, compared with 41 of 301 (13.62%) strict-intergenic events in the ≥6,000-read zone. More strikingly, 260 of the 3,487 low-support events (7.46%) overlapped one of the eight young A/Gf/Tf subfamilies, whereas none of the 301 ≥6,000-read strict-intergenic events did so. The 41 high-support L1-overlapping events instead mapped predominantly to other, generally older subfamilies. Thus, enrichment of evolutionarily young L1 loci represents a distributed low-abundance tumor-associated population rather than a component of the strongly expanded integration class.

At the individual-tumor level, young A/Gf/Tf-associated integrations were heterogeneously represented but remained concentrated predominantly within the low-support compartment (**Fig. 2H**). Their tumor-to-tumor variation was not explained simply by total integration burden (Supplementary material 1). Importantly, high-support integrations were readily detected in other L1 subfamilies within the same tumors, demonstrating that the low-support distribution of young A/Gf/Tf-associated events was not a general property of L1-targeted integrations.

Additional analyses did not support genomic abundance or basal hepatocyte transcription as sufficient explanations for the subfamily redistribution (Supplementary material 1: **Fig. S3A**-**B**). Notably, L1MdTf_III showed one of the strongest reservoir-to-tumor shifts despite a relatively limited genomic TA target space and low basal hepatocyte expression, whereas L1Lx_IV was depleted despite measurable hepatocyte expression and a closely comparable genomic TA target space (∼597,000 versus 584,424 sites). Thus, neither the number of potentially targetable genomic sites nor basal transcription was sufficient to explain the selective representation of particular L1 subfamilies in tumors.

Together, these analyses reveal two readily distinguishable components selected from the much larger integration landscape generated by somatic 3×fOnc3 mutagenesis: a high-support compartment enriched for integrations affecting protein-coding genes and a broad low-support compartment containing a disproportionate representation of integrations within selected L1 subfamilies. The latter pattern is most consistent with tumor-stage selection or retention associated with specific L1 genomic contexts, although its underlying functional mechanism remains unresolved.

Of the final 22,193 animal-level 3×fOnc3 integration events, 14,278 overlapped at least one Ensembl release 116 gene, yielding 16,350 event–gene assignments across 9,644 unique hit genes. We next used this genic integration landscape to identify and mechanistically classify candidate drivers selected during liver tumorigenesis.

### SB–Fah-adjusted recurrence prioritizes candidate driver genes

The enrichment of protein-coding integrations within the high-support tumor compartment suggested that a substantial component of strong clonal selection acted through protein-coding loci. However, because lower-support protein-coding integrations may also represent positively selected events, read support was not used for driver-gene identification. Instead, we compared the observed 3×fOnc3 integration burden of each gene with its expected background targeting frequency in the SB–Fah reference dataset.

The suitability of SB–Fah as a gene-level background was supported by the extensive overlap between the two integration landscapes. Among 5,769 canonical protein-coding genes hit by 3×fOnc3, 5,737 (99.45%) were also represented in the SB–Fah dataset, leaving only 32 genes detected exclusively in the 3×fOnc3 panel (Supplementary material 1: **Fig. S4**). Thus, the SB–Fah reservoir provided empirical targeting information for essentially the entire protein-coding gene space sampled by 3×fOnc3 in vivo.

To correct recurrence for this non-uniform background targetability, we generated 10,000 artificial SB–Fah panels matched to the size and animal-level structure of the 3×fOnc3 dataset (for details, see the Methods section). For each gene, we defined recurrence excess as ΔRV₍_g_₎ = RV₍_3×fOnc3,g_₎ − mean(RV₍_Fah,g_₎), such that positive values indicate more frequent targeting in the tumor-selected panel than expected from the empirical SB–Fah background. ΔRV formed a continuous distribution rather than a discrete driver–passenger boundary; we therefore used it as a prioritization measure and selected the 300 genes with the largest recurrence excesses as the primary candidate set (ΔRV top-300; Supplementary Data D2).

We next exploited the three-frame architecture of 3×fOnc3 by identifying frame- and orientation-compatible truncated-protein-overexpression (TPO) configurations. Of 8,083 canonical coding-intron event–gene assignments, 1,647 (20.38%) were TPO compatible. Comparison with matched artificial SB–Fah panels showed that the number of genes containing at least one TPO-compatible event was similar between the observed and control panels (1,334 versus a Fah mean of 1,308; P = 0.223). Instead, TPO-compatible configurations became concentrated within selected genes: their fraction among canonical coding-intron assignments increased from an expected 17.69% to 20.38% (P = 0.000999), while the number of TPO-dominant genes increased from a Fah mean of 10.9 to 26 (P = 0.000999; Supplementary material 1: **Fig. S5**). Thus, tumor evolution preferentially enriched frame- and orientation-compatible configurations within a subset of recurrently targeted genes rather than simply increasing the number of genes accessible to TPO. TPO-compatible integrations spanned the full read-support range across the 16 individually analysed tumors and were not confined to strongly expanded clones (**Fig. 3A**). Nevertheless, every tumor contained at least one TPO-compatible integration in the ≥6,000-read compartment.

**Figure 3.**
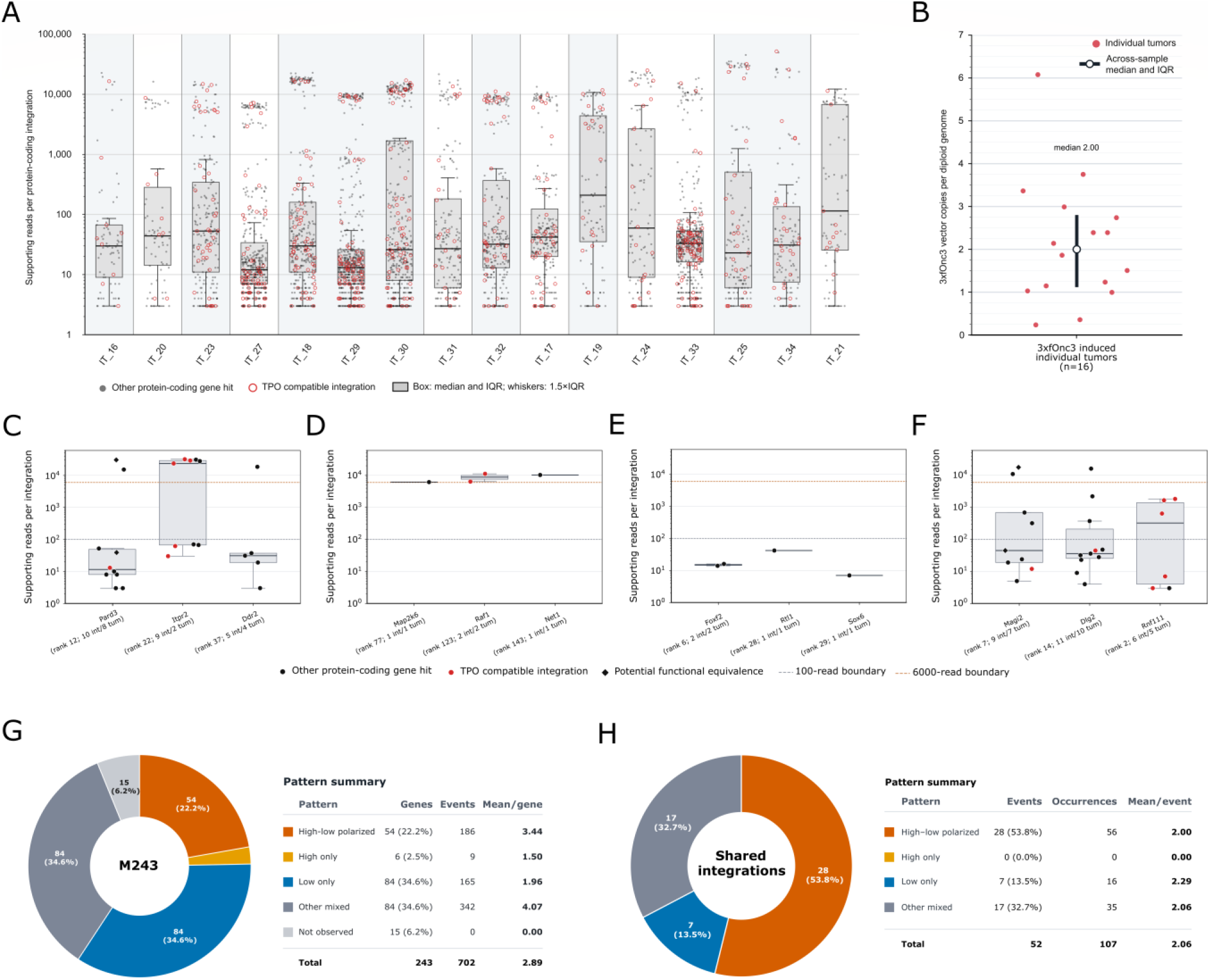
Multiclonality and context-dependent clonal representation of protein-coding driver integrations in 3×fOnc3-induced tumors. (A) Supporting-read distributions of protein-coding gene-associated integrations in the 16 individually dissected 3×fOnc3 tumors. Gray points indicate other protein-coding gene hits and open red circles indicate TPO-compatible integrations. Boxplots show the median and interquartile range (IQR), with whiskers extending to 1.5×IQR. Individual tumors are shown separately; the y-axis is logarithmic. (B) Total 3×fOnc3 transposon copy number measured by qPCR in the same 16 individual tumors and expressed as estimated copies per diploid genome. Red points represent individual tumors; the open symbol and error bars indicate the across-sample median and IQR. The median was 2.00 copies per diploid genome. (C–F) Representative M243 candidate genes illustrating operational read-support patterns across the 16 individual tumors: **(C)** high–low polarized (*Pard3, Itpr2,* and *Ddr2*), **(D)** high only (*Map2k6, Raf1,* and *Net1*), **(E)** low only (*Foxf2, Rtl1,* and *Sox6*), and **(F)** other mixed (*Magi2, Dlg2,* and *Rnf111*). Black points indicate other protein-coding gene hits and red points indicate TPO-compatible integrations. Black diamonds mark potentially functionally equivalent integrations affecting the same intron in the same orientation but at distinct genomic TA target sites; they therefore do not represent identical integration events. Horizontal dashed lines indicate the 100-read and 6,000-read boundaries used for operational classification. Gene rank in the Top315 set and the number of integrations detected among individual tumors are indicated below each gene. Boxplots show the median and IQR, with whiskers extending to 1.5×IQR. The y-axis is logarithmic. (G) Distribution of read-support patterns across the 243 protein-coding genes of the M243 candidate set using integrations recovered from the 16 individual tumors. A gene was classified as high–low polarized when it contained at least one low-support (<100 reads) and at least one high-support (≥6,000 reads) integration and no intermediate-support event; high only and low only genes contained exclusively high- or low-support events, respectively. Any gene containing at least one intermediate-support event (100–5,999 reads) was assigned to the other mixed category. Genes selected into M243 from pooled-tumor or whole-liver libraries but not detected in the individual-tumor panel were classified as not observed. The adjacent table gives the number and percentage of genes in each category, the corresponding number of integration events, and the mean number of events per gene. (H) Read-support patterns of exact integration events shared between spatially distinct individual tumors isolated from the same animal. Shared integrations were required to have the same genomic target position, genomic-flank orientation and dominant 3×fOnc3 construct profile. Fifty-two exact shared integrations were retained, corresponding to 107 tumor-level occurrences. Pattern categories were assigned using the same <100-read, 100–5,999-read and ≥6,000-read boundaries as in panel G. Twenty-eight shared integrations (53.8%) showed a high–low polarized pattern, seven (13.5%) were low only, 17 (32.7%) showed other mixed patterns, and none was high only. The adjacent table shows the number of shared integration events, their tumor-level occurrences and the mean number of occurrences per shared event.

Gene-specific TPO enrichment was therefore used as a complementary driver rescue axis (for details, see the Methods section). Twenty-five genes fulfilled the TPO enrichment rescue criterion; 10 were already present in the ΔRV top-300, whereas 15 additional genes were recovered exclusively through TPO enrichment, producing a final Top315 candidate-driver set (Supplementary Data D2). Restricting Top315 to protein-coding genes yielded the M243 driver candidate set (Supplementary Data D2), of which 123 genes (50.6%) contained at least one TPO-compatible integration and 25 (10.3%) were TPO dominant. A sex-stratified sensitivity analysis identified nine Top315 driver candidates detected exclusively in males and one exclusively in females; the complete analysis is provided in Supplementary material 1.

The large number of integrations identified in individual tumors suggested a high degree of multiclonality. To determine whether this pattern could instead reflect unusually high transposon copy numbers within a small number of dominant clones, we measured total 3×fOnc3 copy number by qPCR. The mean vector burden was 2.14 ± 1.47 copies per diploid genome (n = 16; median, 2.00), comparable to the previously measured 1.83 copies per diploid genome in the SB–Fah system (**Fig. 3B**). In contrast, individual tumors contained a median of 534 distinct integrations, including a median of 216.5 protein-coding integrations (**Fig. 2A and C**). This combination of low vector copy number and hundreds of distinct integration sites demonstrates extensive multiclonality within the analysed tumors.

We next examined whether strong clonal representation was a reproducible property of individual candidate driver genes. Using operational low (<100 reads), intermediate (100–5,999 reads) and high (≥6,000 reads) support zones, M243 genes were classified as high–low polarized, high only, low only or other mixed (**Fig. 3C–G**). Fifty-four genes (22.2%) showed a high–low polarized pattern, whereas only six (2.5%) were represented exclusively by high-support integrations; 84 were low only, 84 showed mixed patterns and 15 were not represented among the individually analysed tumors. Heterogeneous support states became increasingly apparent among repeatedly targeted genes, indicating that high–low polarization reflects repeated sampling of recurrently selected genes rather than a discrete biological driver class. Most importantly, recurrent targeting of the same candidate gene did not imply reproducibly strong clonal expansion. Integrations affecting individual genes could be represented by fewer than 100 reads in some tumors and more than 6,000 in others, in several cases spanning more than three orders of magnitude (**Fig. 3C**). Because different integrations within the same gene may differ in position, orientation or functional consequence, gene-level comparisons alone cannot distinguish mutation-specific effects from differences in the genetic, cellular or microenvironmental context in which each lineage evolved. Nevertheless, the widespread heterogeneous patterns show that realized clonal representation is not determined by driver-gene identity alone.

We therefore asked whether such divergence persisted when the integration event itself—not merely the affected gene—was held constant. Fifty-two exact integrations, defined by identical genomic target position, genomic-flank orientation and dominant 3×fOnc3 construct profile, were recovered from more than one spatially distinct tumor within the same animal, corresponding to 107 tumor-level occurrences (**Fig. 3H**; Supplementary material 1: **Fig. S6**). Of these shared integrations, 28 (53.8%) showed a high–low polarized pattern, being represented by <100 reads in at least one tumor but ≥6,000 reads in another; none was observed exclusively in the high-support state. Seven were low only and 17 showed mixed support patterns. Thus, strongly divergent clonal representation was frequent even when genomic position, integration orientation and transposon construct were identical.

Because exact shared integrations strongly indicate common lineage ancestry, their recovery from spatially distinct tumors also supports substantial intrahepatic dispersal of integration-bearing lineages. The markedly different representation of the same lineage across separate lesions demonstrates that the extent of clonal expansion is strongly dependent on the tumor context in which that lineage is represented.

Together, these analyses identify a recurrence- and TPO-enriched set of candidate liver-cancer drivers while revealing extensive multiclonality and strong context dependence of their clonal representation. The resulting M243 protein-coding candidate set provides a genetically and mechanistically prioritized basis for examining cross-species cancer relevance in the human genome.

### Human orthologs of 3×fOnc3 driver candidates are enriched for cancer-associated mutations and pathways

To examine the relevance of the mouse driver landscape to human cancer, we mapped the M243 protein-coding candidate set to human orthologs. Of the 243 mouse genes, 228 could be assigned a single recommended human ortholog and constituted the H228 panel used for cross-species analyses. Eighteen of these genes (7.89%) were represented in the COSMIC Cancer Gene Census (17 Tier 1 and one Tier 2). In 10,000 matched human gene sets, the mean number of CGC genes was 8.82, corresponding to a 2.04-fold enrichment in H228 (empirical P = 0.0033). Thus, the mouse-derived candidate set was significantly enriched for independently established human cancer genes.

We next asked whether the human orthologs showed broader evidence of cancer-associated somatic mutation. Protein-coding mutations were analysed across 10,443 TCGA PanCancer Atlas tumors from 32 studies using the MC3 dataset. After exclusion of CCDC162P, which lacked an available MANE protein length, the resulting H227 panel was compared with 10,000 protein-length-matched random gene sets. Median per-gene mutation prevalence was significantly elevated for nonsynonymous mutations (BH-adjusted empirical q = 3.3 × 10⁻⁴), missense mutations (q = 3.3 × 10⁻⁴) and truncating mutations (q = 0.002), whereas synonymous mutation prevalence was not significantly enriched (q = 0.145) (**Fig. 4A**). The selective enrichment of protein-altering, but not synonymous, mutations argues against a nonspecific increase in mutation prevalence across the candidate panel.

**Figure 4.**
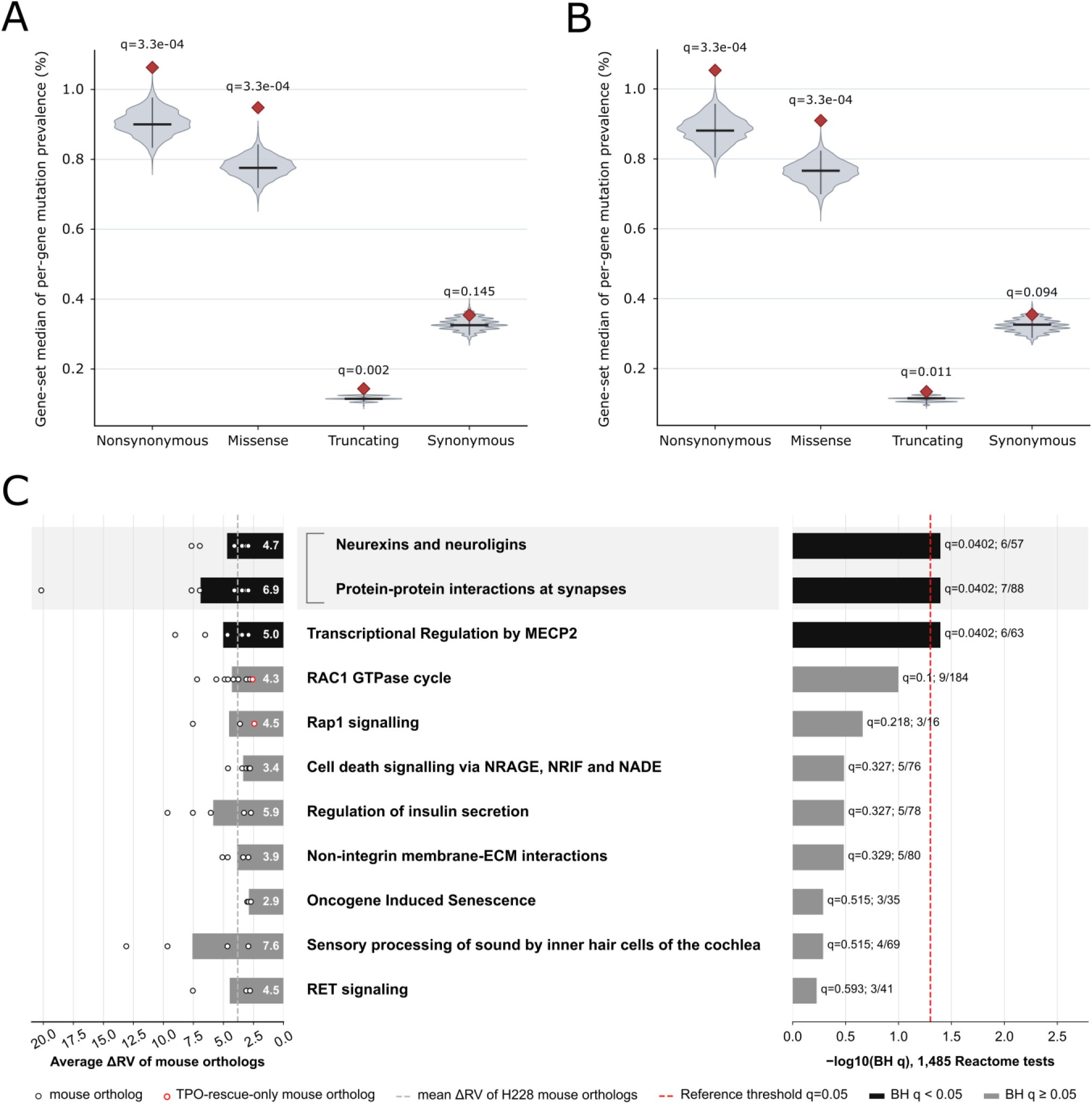
Human orthologs of 3×fOnc3 driver candidates are enriched for cancer-associated mutations and pathways. (A) Pan-cancer mutation prevalence of the 227 human protein-coding orthologs with MANE-defined protein lengths (H227). Red diamonds indicate the observed median per-gene prevalence of nonsynonymous, missense, truncating, or synonymous mutations across 10,443 TCGA PanCancer Atlas tumors. Violin plots show the corresponding null distributions obtained from 10,000 protein-length-matched random gene sets. Horizontal black lines indicate null-distribution medians. Empirical *q* values are shown above each mutation class. (B) The same analysis after removal of the 18 Cancer Gene Census (CGC) genes from the candidate set and matched background pool (H209). Enrichment remained significant for nonsynonymous, missense, and truncating mutations, whereas synonymous mutations were not enriched. (C) Reactome pathway enrichment of the 228 high-confidence human orthologs (H228). Eleven representative nominally enriched pathway groups are shown after redundancy collapsing. The left panel displays ΔRV values of the corresponding mouse orthologs; open circles denote individual genes, red-outlined circles indicate candidates added exclusively through TPO-enrichment rescue, and horizontal bars show the mean ΔRV for each pathway. The grey dashed line marks the mean ΔRV across all H228 mouse orthologs. Numbers within bars indicate pathway-specific mean ΔRV values. The right panel shows −log10(BH-adjusted *q*) for the corresponding Reactome terms; black bars denote BH *q* < 0.05 and grey bars BH *q* ≥ 0.05. The red dashed line indicates *q* = 0.05. Labels report BH-adjusted *q* values and the number of candidate genes overlapping the pathway relative to total pathway size. The two synaptic terms grouped by the bracket belong to the same redundancy cluster and are displayed separately because they represent closely related but distinct Reactome annotations. Reactome enrichment was tested by one-sided hypergeometric analysis across 1,485 pathways, followed by Benjamini–Hochberg correction.

To determine whether this signal was driven primarily by the 18 established CGC genes, we repeated the analysis after removing CGC members from both the candidate panel and matched background. The resulting H209 non-CGC panel retained significant enrichment for nonsynonymous and missense mutations (q = 3.3 × 10⁻⁴ for both) and truncating mutations (q = 0.011), whereas synonymous mutations again remained non-significant (q = 0.094) (**Fig. 4B**). Thus, the pan-cancer mutation enrichment is not driven by already established CGC genes and extends the cross-species cancer relevance of the candidate set to a substantially larger group of currently non-CGC genes.

Reactome analysis of H228 identified three significantly enriched terms representing two principal functional themes (**Fig. 4C**). Neurexins and neuroligins and the closely related Protein-protein interactions at synapses pathway formed a shared synaptic cell-adhesion/protein-interaction cluster, whereas Transcriptional Regulation by MECP2 defined a second theme centred on transcriptional regulation. Additional nominally enriched pathways were displayed after redundancy reduction together with ΔRV values of the corresponding mouse orthologs. Nine of the eleven displayed functional groups showed a mean ΔRV above the H228-wide mean, indicating that the pathway-level signals were generally supported by recurrently targeted genes in the mouse screen.

Among the nominally enriched pathways, the RAC1 GTPase cycle was notable for the mechanistic coherence of its candidate members, several of which regulate or transmit Rac/Rho-family signalling. FARP1, one of these genes, entered the final candidate set through the independent TPO-enrichment rescue criterion. Although the RAC1 pathway did not remain significant after multiple-testing correction, its convergence with independently prioritized 3×fOnc3 candidates identifies it as a biologically suggestive component of the broader driver landscape.

## Discussion

The 3×fOnc3 system was designed to mitigate integration-pattern distortions characteristic of traditional SB-based cancer-gene discovery screens, broaden access to promoter-driven truncated-protein configurations, and separate the generation of somatic mutations from their subsequent selection during tumor evolution. Hydrodynamic delivery of the 3×fOnc3 transposons together with SB100 produces a temporally restricted burst of somatic integration, with detectable transposition largely ceasing within approximately one week [7]. The subsequent ∼12-month interval therefore primarily reflects prolonged in vivo selection rather than continuous mutagenesis, reducing late mutational input while allowing an initially broad integration landscape to undergo extended selection. Another key feature is the use of the SB–Fah dataset as an empirical hepatocyte integration reference. Unlike theoretical genomic TA-site space, this reservoir incorporates hepatocyte-specific genomic accessibility and intrinsic SB targeting preferences under closely matched somatic delivery conditions. SB–Fah is not a neutral genomic null, because corrected hepatocytes undergo Fah-driven regenerative expansion, but its near-complete overlap with the protein-coding target space sampled by 3×fOnc3 makes it a biologically relevant reference for background targetability. Its compact read-support distribution further supports treating SBE-seq read support as a semiquantitative measure of clonal representation without equating read counts with absolute cell numbers.

Tumor selection reshaped this integration landscape in a non-uniform manner. Protein-coding integrations were preferentially represented within the high-support compartment, whereas the broader low-support population was relatively enriched for repetitive genomic regions. These zones should not be interpreted as driver and passenger classes: low-support protein-coding integrations can represent selected driver events, and read support reports realized clonal expansion rather than intrinsic oncogenic potential alone. The data instead support a continuum of integration-bearing lineages experiencing heterogeneous selective growth outcomes.

A particularly unexpected component of this landscape was the selective overrepresentation of integrations within endogenous L1 elements. Enrichment was strongly subfamily dependent and included several evolutionarily young A, Gf and Tf subfamilies, whereas L1Lx_IV showed the opposite trend. Neither genomic TA-site availability nor basal hepatocyte transcription reproduced this pattern. The opposing behaviour of L1MdTf_III and L1Lx_IV despite their closely comparable genomic TA target spaces is particularly informative, arguing against simple preferential accessibility of abundant or transcriptionally active L1 sequences. Importantly, integrations within the young A/Gf/Tf group were confined to the low-support compartment, although high-support events were readily detected in other L1 subfamilies. Their distribution is compatible with selection or retention associated with particular young-L1 genomic contexts without requiring a strong cell-autonomous proliferative effect. Whether these integrations are themselves functional or instead mark genomic or cellular states linked to another selectable phenotype will require dedicated mechanistic experiments. Notably, recent studies show that young L1MdTf subfamily elements possess cis-regulatory potential, influencing the expression of genes in their vicinity [11].

Gene-level prioritization used excess recurrence relative to the empirical SB–Fah background rather than raw integration frequency. This is important because gene size, TA-site content, chromatin accessibility and other locus-specific properties can strongly influence the probability of recovering an SB integration independently of tumor selection. ΔRV therefore provides a background-adjusted ranking of recurrently selected genes, but not a binary driver classifier. Its continuous distribution and the operational nature of the top-300 cutoff mean that the final Top315 set should be viewed as a prioritization of the strongest recurrence- and TPO-supported candidates.

The second prioritization axis exploited the established ability of Onc-family promoter–splice-donor cassettes to generate downstream fusion transcripts and truncated or ectopically expressed products. The distinctive contribution of 3×fOnc3 is systematic provision of all three translational reading frames, thereby broadening the intronic configurations capable of generating TPO-compatible downstream products. Tumor selection did not simply increase the number of genes containing a TPO-compatible event; instead, frame- and orientation-compatible configurations became concentrated within selected genes. Gene-specific TPO enrichment consequently provided an orthogonal rescue axis for candidates that ranked below the strongest ΔRV signals. The importance of three-frame access was also evident in the selected landscape: among the 25 TPO-dominant protein-coding candidates, the historical Onc3 configuration would have provided access to at least one dominant TPO intron in only five, and none of the seven genes showing complete intron-level convergence could reproduce the observed configuration with the historical construct. Thus, the three-frame design materially expands access to selectable protein-altering configurations.

An equally important result is that driver identity and clonal dominance were not equivalent. Individual candidate genes were represented across widely different read-support states, indicating that recurrent targeting of the same gene does not prescribe a reproducible final clone size. At the gene level, such heterogeneity could still reflect integration-specific functional differences or additional alterations acquired by individual lineages. The shared-integration analysis substantially narrows these alternatives. Exact integrations with the same genomic position, orientation and construct identity were recovered from spatially distinct tumors within the same animal, and more than half displayed strongly divergent high–low representation. Because such events strongly indicate common lineage ancestry, the same integration-defined lineage can remain minor in one lesion while becoming strongly expanded in another. Branch-specific early integrations or later endogenous alterations may modify individual lineages, but are unlikely to provide a general explanation for such a widespread pattern. Realized clonal expansion is therefore strongly dependent on the genetic, cellular and/or local microenvironmental context in which a lineage evolves.

The recovery of multiple shared integration-defined lineages from spatially separated tumors also supports substantial intrahepatic lineage dispersal. This interpretation is consistent with lineage-tracing and imaging studies showing that hepatocyte populations can expand across lobular boundaries and undergo spatial redistribution or collective migration during liver regeneration [12, 13]. These studies demonstrate that substantial hepatocyte movement is biologically plausible.

Together with the low vector copy number and hundreds of integrations recovered from individual tumors, the shared-lineage data support a model in which multiple mutagenized hepatocyte lineages can contribute to developing tumor sites and subsequently undergo different degrees of local expansion, rather than tumor growth being reducible to uniform expansion of a single dominant clone. Consistent with this, increasing evidence has been found in recent years demonstrating the polyclonal origin of solid tumors [14, 15].

The young-L1-associated population is compatible with this broader multiclonal picture. Its selective enrichment without emergence into the high-support compartment suggests that tumor evolution can retain integration-bearing populations whose contribution, if functional, need not involve clonal dominance. This raises the possibility of distributed or context-dependent roles within a multiclonal tumor environment. This selective enrichment of young L1 hits may help the developing tumor acquire a particular cancer hallmark capability [16]. While in the multiclonal tumor microenvironment, where cooperative interactions among subclones occur in line with the public goods theory [17], particular clones do not need to possess all hallmark capabilities—and may not even be exceptionally proliferative—their presence in the tumor can still be beneficial.

The human analyses provide an independent cross-species test of the candidate landscape. Mouse genes were prioritized without using human cancer annotations, yet their human orthologs were enriched for established Cancer Gene Census genes and for protein-altering somatic mutations across TCGA PanCancer Atlas tumors. Synonymous mutations showed no corresponding enrichment, arguing against a nonspecific increase in baseline mutability. Importantly, the protein-altering mutation signal persisted after established CGC genes were removed, extending the cross-species cancer relevance of the candidate set to a substantially larger group of currently non-CGC genes. These pan-cancer analyses test general cancer relevance rather than HCC specificity, and they do not establish driver activity for every individual M243 gene.

Pathway analysis provided a complementary level of convergence. The significant Reactome terms grouped into a synaptic cell-adhesion/protein-interaction theme and a transcriptional-regulation theme. Although the terminology of the former derives from neuronal biology, genes such as SHANK2 and EPB41L5 have established roles in cancer-associated signalling, adhesion and invasion, making convergence on scaffold and cell-interaction machinery a more plausible interpretation than acquisition of a neuronal phenotype [18, 19].

The Transcriptional Regulation by MECP2 term included MET, which is particularly informative because it was also the only overlap between our prioritized candidate set and the 19 common insertion sites reported in the earlier T2/Onc3 liver screen [6]. Its recovery provides a point of continuity with the earlier platform while the substantially broader 3×fOnc3 candidate landscape reflects expanded access to selectable driver configurations. The pathway itself should nevertheless be interpreted as a broader transcriptional-regulatory module rather than as evidence for a specific MECP2-dependent mechanism in these tumors.

The nominally enriched RAC1 GTPase cycle provides a further mechanistically coherent example, although it did not remain significant after multiple-testing correction. Its members included several Rac/Rho-family regulators, and FARP1 is notable because it entered the final candidate set through the independent TPO-enrichment rescue axis rather than the ΔRV top-300. FARP1 has also been identified experimentally as a Rac1 GEF mediating receptor-tyrosine-kinase-dependent cancer-cell motility, including downstream of c-MET [20]. Such convergence illustrates how mechanistically distinct prioritization layers can identify biologically coherent candidates that would be missed by recurrence ranking alone, while the nominal statistical status of the pathway warrants cautious interpretation.

Together, these findings position 3×fOnc3 as a background-aware somatic mutagenesis platform that links driver prioritization with clonal tumor evolution. Cross-species enrichment for established cancer genes, protein-altering human mutations and defined functional pathways further supports the cancer relevance of the resulting candidate landscape while leaving a substantial non-CGC component for future investigation. In the future, this technological platform could be suitable for analyzing the relationship between HCC driver selection and genetic background, as well as the effects of specific conditions such as fatty liver disease, certain infections, and drugs. Furthermore, with the help of modern nucleic acid packaging and targeting technologies, the system could be adapted beyond the organ-specific environment examined here, thereby enabling the study of additional cancer types.

## Methods

### Animal care and maintenance

C57BL/6J mice were obtained from The Jackson Laboratory and bred and maintained in the Central Animal House at the Biological Research Centre (Szeged, Hungary). Specific pathogen-free status was confirmed quarterly according to FELASA (Federation of European Laboratory Animal Science Associations) recommendations (FELASA 2014, doi: 10.1177/0023677213516312). All animal experiments were conducted according to protocols approved by the Institutional Animal Care and Use Committee of the Biological Research Centre under the supervision of the Governmental Office for Csongrád County, Directorate of Food Chain Safety and Animal Health. The approval number was XVI./801/2018. Chapter 1 Methods

### Plasmid construction

The pT2/Onc3-F0, pT2/Onc3-F1 and pT2/Onc3-F2 mutagenic transposon plasmids were derived from pT2/Onc3 [9]. Briefly, the QuikChange Site-Directed Mutagenesis protocol (Agilent) was used to introduce Kozak–ATG elements providing the corresponding translational reading frames upstream of the splice-acceptor element of pT2/Onc3. Distinct two-nucleotide barcodes were subsequently introduced into the left SB inverted terminal repeats (ITRs) of pT2/Onc3-F1 and pT2/Onc3-F2 using the same approach, whereas the left ITR of pT2/Onc3-F0 retained the wild-type sequence. Oligonucleotides used for plasmid construction are listed in Supplementary Data D1. The same two-nucleotide barcode substitutions were introduced into the pT2/SVNeo transposon vector (Addgene #26553) [21] to generate pT2/SVNeo-F1BC and pT2/SVNeo-F2BC for comparative transposition-efficiency testing in mammalian cells.

### Transposition assay

HepG2 cells were purchased from ATCC and cultured in Dulbecco’s modified Eagle’s medium (DMEM) (Biosera) supplemented with 10% fetal bovine serum (FBS; Gibco) and 1% penicillin-streptomycin (P-S; HyClone) in the presence of 5% CO2 at 37 °C. Cells were transfected with 50 ng of either pT2/SVNeo (wt, F0BC) or its barcoded variants pT2-SVNeo-F1BC or pT2-SVNeo-F2BC along with 50 ng of the pCMV/SB10 (Addgene #24551) [22] transposase helper plasmid —with pUC19 added to a total DNA amount of 500 ng per transfection—using FuGENE® HD transfection reagent (Promega) according to manufacturer’s instructions. Selection of stable transfected cells was performed using neomycin (G418; Biosera). After 14 days of selection colonies of cells were stained with 0.6% methylene blue, air-dried, and counted.

### Nucleic acid isolation

For SBE-seq and transposon copy-number assessment by qPCR, whole-liver or tumor samples were lysed in 100 mM Tris-HCl (pH 8.0), 5 mM EDTA (pH 8.0), 200 mM NaCl and 0.2% SDS and incubated overnight at 50°C in the presence of 300 μg/mL Proteinase K (VWR Chemicals). One milliliter of lysate was subjected to standard phenol/chloroform extraction. DNA was precipitated with three volumes of absolute ethanol in the presence of 0.1 volume of 3 M sodium acetate. DNA pellets were washed with 70% ethanol, air-dried and dissolved in nuclease-free water (Sigma-Aldrich). DNA concentration was determined using a Qubit 3.0 Fluorometer (Thermo Fisher Scientific).

### Hydrodynamic tail-vein injection

Plasmids used for hydrodynamic tail-vein injection were prepared using the NucleoBond Xtra Maxi Plus EF Kit (Macherey-Nagel) and diluted in Ringer’s solution immediately before administration. A volume equivalent to 10% of body weight was injected through the lateral tail vein within 5–8 s into 8-week-old mice, as described previously [23]. Thirty C57BL/6J mice (15 males and 15 females) received the 3×fOnc3 mutagenic construct mixture. The transposon component comprised 40 μg total plasmid DNA: 18 μg pT2/Onc3-F0, 12 μg pT2/Onc3-F1 and 10 μg pT2/Onc3-F2, corresponding to 45%, 30% and 25% of the transposon mixture, respectively. These proportions were selected to approximate the unequal representation of the corresponding intron phases in the mouse genome [10]. The transposon mixture was co-administered with 4 μg of the pcGlobin2-SB100 [24] transposase helper plasmid. Animals were sacrificed approximately 12 months after injection.

### Sequencing of transposon insertion sites using SBE-seq

SB transposon left-ITR/genome junctions were sequenced using SBE-seq as described previously [7]. SBE-seq library preparation and Illumina sequencing used the same previously published parameters as those applied to the SB–Fah dataset.

Three Illumina MiSeq runs covered the C57BL/6J (B6) 3×fOnc3 experimental panel, which comprised 32 sequenced libraries from 18 animals: 17 individually dissected tumor libraries, 11 pooled-tumor libraries and four whole-liver libraries. No integration call passed final quality control in either the pooled-tumor library or the individually analysed tumor library derived from one animal. Consequently, the downstream B6 dataset contained 30 non-empty libraries from 17 animals: 16 individual tumors, 10 pooled-tumor libraries and four whole-liver samples.

### Re-analysis of the SB–Fah reference integration dataset

The previously published SB–Fah therapeutic gene-transfer dataset [7] was used as an empirical SB integration reference in mouse hepatocytes. To ensure direct analytical comparability, SB–Fah transposon integrations were re-identified and processed using the same integration-site calling framework applied to the B6 3×fOnc3 dataset.

### Integration-site calling using complementary mapping branches

Sleeping Beauty transposon–genome junctions were identified from Read 2 sequences, and extracted genomic flanks were mapped to the GRCm39 primary assembly using two complementary approaches.

In the strict alignment branch, complete genomic flanks were aligned using BWA-MEM v0.7.17 and retained only when the primary alignment was uniquely supported (MAPQ ≥60, no XA or secondary-alignment evidence, and AS−XS ≥10 when an alternative alignment score was present). At least 16 uninterrupted and exactly matching genomic bases were required immediately adjacent to the transposon junction; mismatches, insertions/deletions and clipping were not permitted within this protected junction-proximal anchor.

In parallel, every extracted genomic flank was analysed using an all-read, junction-anchored variable-length exact-prefix branch. Flanks were queried with BWA v0.7.17 fastmap using a minimum SMEM length of 16 bp (bwa fastmap -l 16). From the resulting exact matches, the custom caller selected the longest exact reference match beginning at the first genomic base adjacent to the transposon junction and retained the read only when this prefix occurred uniquely in the complete reference assembly. This variable-length exact-prefix procedure allowed junctions to be assigned when sequencing errors, adapter read-through or ligation-derived sequence affected the distal portion of the genomic flank.

The two mapping branches were integrated by read identifier. Concordant assignments of the same read to the same genomic coordinate were counted once, whereas reads assigned to conflicting coordinates by the two branches were excluded. Branch provenance was retained for downstream quality control. The same dual-mapping and conflict-resolution procedure was applied to the SB–Fah and 3×fOnc3 panels.

Calls mapping outside the predefined set of eligible main contigs were excluded before downstream correction and filtering.

For the 3×fOnc3 panel, construct frame (F0/F1/F2), transposon orientation and supporting-read count were recorded for each integration site. The transposed construct variant was identified through its two-nucleotide barcode located in the left SB ITR within the SBE-seq amplicon (Supplementary Data D1). Because the sequenced left-ITR genomic flank is oriented opposite to the internal CAG promoter of the Onc3 construct, promoter–gene orientation was classified as sense when the genomic-flank strand and gene strand were opposite and as antisense when they were identical.

### TA-stutter/PCR-slippage correction

Integration calls were audited for local families of closely spaced canonical TA target sites occurring within the same biological library, reference contig and genomic-flank orientation. These families typically formed 2- or 4-bp repeat ladders and consisted of low-support satellite calls surrounding a dominant coordinate. We refer to this pattern as *TA stutter*. Local sequence periodicity, read-support structure and, for 3×fOnc3 integrations, concordant F0/F1/F2 construct profiles were consistent with positional call inflation caused by PCR slippage or a related amplification/calling artefact.

TA-stutter-compatible families were identified from coordinate spacing, local reference-sequence periodicity or TA-alternating low complexity, read-support structure and, where available, construct-profile concordance. Each accepted family was collapsed to its highest-support coordinate, and supporting reads from all family members were transferred to that representative, thereby preserving total read support.

In the B6 3×fOnc3 panel, TA-stutter correction reduced the dataset from 26,627 to 26,013 sample-level calls by merging 614 satellite calls in 314 families, while preserving all 21,371,403 supporting reads. In the SB–Fah reference, the corresponding correction reduced 911,080 to 908,273 event rows by merging 2,807 satellite calls in 916 families, again without changing total read support.

### Run-specific filtering of cross-sample technical echo in the 3×fOnc3 panel

Within each MiSeq run, exact integration events shared between separately indexed libraries were examined for evidence of cross-sample technical echo or carryover. Events observed in different MiSeq runs were not subjected to echo filtering.

An exact integration event was defined by genomic target interval, genomic-flank orientation and dominant F0/F1/F2 construct profile. Calls sharing the same genomic target and flank orientation but carrying different construct-frame profiles were therefore treated as distinct integration events and retained.

Echo classification was applied to identical events within the same MiSeq run when the highest-support occurrence contained at least 1,000 reads. At shared sites, the highest-support library assignment was retained when its read support was at least tenfold higher than that of the next-highest library, whereas the lower-support assignments were excluded. When no library showed tenfold dominance, all assignments at that site were retained.

Application of the unified same-run echo filter to the TA-stutter-corrected 3×fOnc3 panel retained 25,203 sample-level calls. One additional low-support call in IT_22 showed an otherwise characteristic cross-sample echo pattern despite originating from a donor event below the operational 1,000-read activation threshold and was therefore excluded after manual review. The resulting echo-corrected B6 dataset contained 25,202 sample-level calls.

No analogous run-specific echo classifier was applied to the SB–Fah reference. Only two SB–Fah liver libraries were multiplexed per MiSeq run, providing substantially less information for reconstructing run-specific donor-to-recipient echo patterns than the more densely multiplexed 3×fOnc3 runs. Moreover, the SB–Fah read-support distribution was narrow: among the 908,273 TA-stutter-corrected SB–Fah calls, median support was 48 reads, mean support was 53.45 ± 35.08 reads, and the maximum was 772 reads; the median of the 12 sample-specific medians was 49.5 reads. Thus, no SB–Fah integration fulfilled the ≥1,000-read high-support donor criterion used to activate the echo classifier.

### Exclusion of the Onc3-homologous region of the mouse *En2* gene

The 431-bp *En2* segment carried by the 3×fOnc3 constructs exactly matches GRCm39 chr5:28374841–28375272. Integration calls mapping within this interval could not be unambiguously distinguished from construct-derived junctions, including potential plasmid-to-plasmid transposition products, and were therefore excluded from downstream analyses.

Exclusion of the construct-homologous *En2* interval removed 96 sample-level calls, reducing the B6 panel from 25,202 to 25,106 sample-level calls. At the animal-event level, this corresponded to removal of 81 events, from 22,274 to 22,193 events. The final B6 integration panel therefore comprised 25,106 sample-level calls corresponding to 22,193 animal-level integration events.

### Animal-level consolidation and final construct-frame assignment

After integration-level corrections were completed, F0/F1/F2 evidence was re-evaluated from the aggregated profile counts associated with each retained sample-level call. At least 80% of reference-matching read support had to carry an assignable construct profile. Calls supported by a single profile were assigned automatically. For mixed-profile calls, automatic assignment required an untied dominant profile accounting for at least 90% of assigned reads and supported by at least 10 reads. Calls with lower dominance, tied profiles or insufficient assigned support were classified as unresolved. Among the 19,936,473 integration-supporting reads retained in the final sample-level callset, the two-base barcode/profile match uniquely resolved an F0, F1 or F2 profile for 19,873,558 reads (99.684%). The remaining 62,915 reads (0.316%) produced an ambiguous best-profile match.

Sample-level calls were subsequently consolidated at the animal-event level. An animal-level event received a final F0, F1 or F2 assignment only when all constituent sample-level calls were automatically callable and supported the same frame. Events containing non-callable or discordant frame evidence were retained in the integration dataset but excluded from automatic frame-dependent downstream classification.

### Ensembl annotation of the final integration set

The final 22,193 animal-level events were annotated against Ensembl release 116 on GRCm39 while retaining overlapping genes. Exact gene-span overlap was defined using Ensembl GTF gene intervals without upstream or downstream extensions. Because a single integration could overlap more than one annotated gene, one event could generate multiple event–gene assignments.

This annotation yielded 14,278 genic events, 16,350 event–gene assignments and 9,644 unique hit genes. The SB–Fah reference dataset was annotated using the same procedure.

### Read-support distribution analysis and definition of the high-support zone

Supporting-read distributions were compared between 12 SB–Fah whole-liver samples, two macroscopically tumor-free 3×fOnc3 whole-liver samples, two tumor-bearing 3×fOnc3 whole-liver samples and 16 non-empty individually dissected 3×fOnc3 tumors.

For visualization of group-level read-support distributions, each biological sample contributed 2,000 deterministic quantiles spanning its within-sample integration-support distribution. This gave each biological sample equal weight irrespective of the number of integration events recovered from that sample.

Inspection of the sample-balanced distributions identified 6,000 supporting reads as the empirical lower boundary of an upper-support component present in individual tumor samples but absent from the SB–Fah reference. The 6,000-read threshold was therefore used to stratify integrations into <6,000- and ≥6,000-read zones; it should be interpreted as an empirical analytical boundary.

To evaluate the robustness of the tumor-specific high-support component, the 16 non-empty individual tumors were compared with the 12 SB–Fah whole-liver samples using sample-level rather than event-level statistics. For each biological sample, the fraction of events supported by ≥6,000 reads, the 99th percentile of read support, the Gini index of the read-support distribution and the fraction of total supporting reads contributed by the upper 1% of events were calculated. Group differences were evaluated using two-sided Mann–Whitney tests. Confidence intervals for differences between group medians were estimated from 10,000 bootstrap resamplings performed at the biological-sample level. Read-support distributions were additionally normalized within each sample to reads per million accepted supporting reads (RPM) to determine whether the high-support tumor component could be explained by differences in sequencing allocation.

### Genomic-context annotation of strict-intergenic integrations

Accepted SB–Fah and 3×fOnc3 integrations that did not overlap any complete Ensembl release 116 gene span were classified as strict-intergenic events. For the analyses shown in **Fig. 2E–G**, 3×fOnc3 events were additionally stratified according to supporting-read count.

Regulatory annotations were obtained from Ensembl Regulation release 116 on GRCm39 (Mus_musculus.GRCm39.regulatory_features.v116.gff3.gz and Mus_musculus.GRCm39.regulatory_activity.v116.tsv.gz). A regulatory feature was designated liver developmental/neonatal active when at least one of the liver E11.5, E12.5, E13.5, E14.5, E15.5, E16.5 or liver 0 d activity fields was annotated as ACTIVE. Integration events were required to overlap the genomic interval of an enhancer, promoter, open-chromatin or CTCF-binding feature directly. This classification was used specifically as an embryonic/neonatal liver regulatory annotation and was not interpreted as evidence of activity in adult hepatocytes. Gene identifiers associated with promoter records in the regulatory GFF were not used for gene assignment because they originated from an earlier Ensembl release; gene spans and TSS positions were instead derived independently from the Ensembl release 116 GTF.

Strict-intergenic events assigned to RepeatMasker sequence were further classified into L1, other LINE, LTR, SINE, DNA transposon, simple/low-complexity sequence, satellite, other repeat or multiple-repeat categories for the analysis shown in **Fig. 2F**.

### L1-subfamily and structural analyses

For L1-specific analyses, accepted integrations were restricted to strict-intergenic events overlapping RepeatMasker L1 annotations. The comparison included 12 SB–Fah whole-liver samples, two macroscopically tumor-free 3×fOnc3 whole-liver samples, two tumor-bearing 3×fOnc3 whole-liver samples and 16 individually dissected 3×fOnc3 tumors. Because all strict-intergenic SB–Fah events included in these analyses had ≤560 supporting reads, direct Fah–3×fOnc3 comparisons of L1 subfamilies, structural properties and orientation were restricted to the <6,000-read stratum.

All RepeatMasker L1 records overlapping an integration were retained in the audit dataset. For analyses requiring a single subfamily assignment, one primary L1 record was selected deterministically by sequentially prioritizing greater event–repeat overlap length, higher RepeatMasker Smith–Waterman score, lower milliDiv, longer genomic repeat span, and finally a stable lexicographic element key defined by contig, coordinates, strand and subfamily. Multiple-overlap information was retained separately, including the complete set of overlapping records and subfamilies and flags for element- and subfamily-level ambiguity.

For subfamily-level comparisons, the contribution of each L1 subfamily was calculated within each biological sample as a fraction of primary L1-overlapping events. Group values shown in **Fig. 2G** represent sample-balanced means of these within-sample fractions. Differences between the 12 SB– Fah whole-liver samples and 16 individual 3×fOnc3 tumors were evaluated using 50,000 sample-label permutations. Two-sided empirical P values were corrected across the tested L1 subfamilies using the Benjamini–Hochberg procedure, and q ≤ 0.05 was considered statistically significant.

A primary RepeatMasker L1 record was classified as near full length when its genomic span was at least 5,500 bp, estimated consensus coverage was at least 90%, and no more than 500 bp was missing from either consensus end. Records of at least 4,000 bp that failed these criteria were classified as long fragments. Genomic sequences corresponding to records of at least 4,000 bp were oriented in the L1 direction and screened for an ATG-to-stop-or-segment-end ORF1-length candidate of 750–1,500 nt followed by an ORF2-length candidate of at least 3,000 nt, separated by −100 to +500 nt. This analysis was used exclusively as a structural length-compatibility screen and was not interpreted as evidence for intact ORFs, transcription, translation or retrotransposition competence.

Sample-balanced structural and consensus-position metrics were compared between individual tumors and SB–Fah samples using 50,000 sample-label permutations with Benjamini–Hochberg correction across four tests.

### Freshly isolated hepatocyte expression annotation of L1 subfamilies

L1-subfamily abundance was estimated from three publicly available CEL-Seq2 libraries generated from freshly isolated wild-type mouse hepatocytes (GSE248554; SRR26941585–SRR26941587). The libraries contained 35-nt single-end 3′ tags. Reads were aligned to GRCm39 using HISAT2 v2.2.1 [25], with splice junctions derived from Ensembl release 116. Up to 100 valid alignments per read were retained to preserve information from reads mapping to multiple repetitive loci. BAM files were sorted by query name using SAMtools v1.13.

TE-subfamily abundance was quantified separately in each biological replicate using TEcount from TEtranscripts v2.2.4 [26], together with the mm39 RepeatMasker transposable-element annotation and Ensembl release 116 gene annotation. TEcount was run in forward-stranded, multimapper-aware mode, with a maximum of 100 iterative allocation steps and a single-end fragment length of 35 nt.

Fractionally allocated TEcount estimates were treated as read-equivalent counts and normalized to the total number of input reads. Although the original CEL-Seq2 libraries contain unique molecular identifiers, UMIs were not used for TE-specific molecule deduplication because multimapping reads were allocated at the alignment level by TEcount. The resulting estimates were therefore interpreted as read-equivalent rather than UMI-defined molecule counts.

### Genomic TA target-space analysis of L1 subfamilies

Genomic TA target space was calculated separately for each L1 subfamily from the UCSC mm39 RepeatMasker table. Records were retained when repeat class was LINE, repeat family was L1, and the repeat name exactly matched the subfamily under analysis. UCSC chromosome identifiers were converted to GRCm39 identifiers, and only contigs represented in the GRCm39 primary-assembly FASTA were retained. Overlapping or directly adjacent RepeatMasker intervals belonging to the same subfamily were merged by genomic union on each contig. GRCm39 sequence corresponding to each interval union was then extracted, and canonical TA dinucleotides were counted only when both bases fell entirely within the union. Thus, a genomic TA site could contribute at most once to the target-space estimate of a given L1 subfamily, irrespective of overlapping RepeatMasker records. Target spaces were calculated independently between subfamilies, such that the same genomic TA site could contribute once to more than one subfamily if it was covered by annotations assigned to different subfamilies.

### Expression annotation of protein-coding genes in the SB-Fah and 3×fOnc3 sample groups

Of the Ensembl 116 protein-coding universe 21,641 genes had a canonical protein-coding transcript. This was used as the primary publication universe. Gene coverage was orientation independent. A gene was classified as Fah+ or 3*×*fOnc3+ when at least one event overlapped its exact gene span, producing four mutually exclusive groups: Fah+/3*×f*Onc3+, Fah+/3*×f*Onc3−, Fah−/3*×f*Onc3+ and Fah−/2*×f*Onc3−. In the canonical 21,641-gene universe, group counts were 5,737 Fah+/3*×f*Onc3+, 12,794 Fah+/3*×f*Onc3−, 32 Fah−/3*×f*Onc3+ and 3,078 Fah−/3*×f*Onc3− genes.

Gene expression was evaluated descriptively across the four Fah/Onc3 coverage groups and was not used for driver selection. Whole-liver expression used an author-supplied TPM matrix for three untreated 7–8-week-old male C57BL/6J liver samples from E-MTAB-6081 (‘199_2_Liver’, ‘199_15_Liver’ and‘199_28_Liver’). Ensembl release 84 matrix rows were linked to the Ensembl 116 gene universe by stable gene identifier. A gene was considered undetected when TPM was exactly zero in all three biological replicates.

Fresh-hepatocyte expression used the three 0-hour libraries in GSE248554 (SRR26941587, SRR26941586 and SRR26941585). Reads were aligned to GRCm39 with HISAT2 v2.2.1 using Ensembl release 116 splice sites, and exonic reads were assigned to Ensembl gene IDs with featureCounts v2.0.3 in forward-stranded mode. Multimapping and ambiguously assigned reads were excluded. Within each sample, reads were collapsed by exact gene ID plus exact 6-base UMI. Detection required at least one uniquely mapped, unambiguously assigned gene–UMI molecule; zero status required zero such molecules in all three replicates.

Genes were classified as undetected when their TPM or exact gene-level UMI count was zero in all three biological replicates. The proportion of undetected genes was reported descriptively for each Fah–Onc3 gene-coverage category in independent whole-liver and freshly isolated hepatocyte datasets.

### Driver gene candidate selection

Our driver gene candidate selection was based on two axes. As the primary axis, we compared the 3xfOnc3 mutation frequency observed in individual genes within the mouse genome to the frequency of Fah therapeutic gene insertions—generated in silico to reflect the same number of integrations and per-animal distribution—as a control. The control dataset derived from our previous SB transposon-based therapeutic gene delivery experiment, in which we identified nearly 1 million SB integrations in mouse hepatocytes [7]. Subsequently, as a complementary axis, we screened for genes exhibiting a high frequency of TPO integrations. The union of these two approaches yielded our narrowed-down list of 315 driver gene candidates.

#### Primary driver gene candidate selection axis

The SB-Fah therapeutic gene-delivery control population comprised 908,273 multiplicity-expanded integration occurrences annotated against Ensembl release 116. Artificial control panels were generated at the event level, before events overlapping multiple genes were expanded into separate event–gene assignments. Each panel reproduced the exact sizes of the 17 non-empty 3×fOnc3 animal bins included in the analysis and therefore contained 22,193 SB-Fah integration occurrences. The fixed bin-size vector was 566, 1,609, 171, 1,157, 2,159, 3,882, 522, 1,536, 2,031, 1,763, 560, 2,458, 401, 417, 2,265, 516 and 180 events.

For each replicate panel, source SB-Fah integration occurrences were sampled without replacement and allocated sequentially to the 17 pseudoanimals in the fixed bin order. Within each pseudoanimal, duplicate oriented loci—defined by chromosome, genomic target interval and genomic-flank orientation—were prohibited. The same oriented locus could nevertheless occur in different pseudoanimals, reproducing the biological possibility of independent integrations at the same locus in different animals. After event sampling, every selected occurrence was assigned to all overlapping Ensembl release 116 genes. For each replicate and gene, the simulated Fah recurrence value was the number of selected events assigned to that gene, whereas pseudoanimal prevalence was the number of pseudoanimals containing at least one event assigned to the gene.

The Fah null distribution comprised 10,000 pseudorandomly sampled panels. For exact computational reproducibility, a master seed of 2026072401 was used to generate a distinct 64-bit seed for each replicate panel. The complete set of panels was used to estimate the gene-specific null distribution of recurrence values. For each gene, the mean recurrence across the 10,000 artificial panels was defined as its expected Fah recurrence value (Fah RV).

The observed 3×fOnc3 recurrence value for a gene was the number of distinct animal-level oriented integration events assigned to that gene. An animal-level oriented event was defined by animal identity, chromosome, genomic target interval and genomic-flank orientation. Consequently, the same oriented locus observed in two different animals contributed two events, whereas repeated detection of the same event in multiple samples from the same animal contributed only once. The same event-counting rule was applied separately within every artificial Fah panel.

The primary effect size, denoted ΔRV, was calculated as: ΔRV₍_g_₎ = RV₍_3×fOnc3,g_₎ − mean(RV₍_Fah,g_₎). A positive ΔRV therefore indicated that a gene received more 3×fOnc3 event–gene assignments than expected from its gene-specific accessibility in an animal-bin- and panel-size-matched SB-Fah therapeutic gene-delivery background. The 300 genes with the highest ΔRV values constituted the primary driver-candidate set (ΔRV top 300) used in subsequent driver analyses.

#### TPO annotation

Canonical coding introns were reconstructed transcript by transcript from the exon and CDS features of Ensembl release 116 protein-coding transcripts; primary analyses were restricted to transcripts tagged as Ensembl_canonical. CDS segments were ordered in transcriptional direction. For each intron, the effective cumulative coding length upstream of the intron was calculated after accounting for the initial Ensembl GTF CDS phase, and the intron frame was defined as its remainder modulo three. Frame consistency was verified against the phase of the downstream CDS segment.

A 3×fOnc3 integration event–gene assignment was classified as TPO compatible only when it (i) overlapped a canonical coding intron of an Ensembl-canonical protein-coding transcript, (ii) placed the CAG promoter in the sense orientation relative to the gene, (iii) had an automatically callable animal-level construct frame and (iv) matched that construct frame to the inferred intron frame. The final 3×fOnc3 panel contained 8,083 canonical coding-intron event–gene assignments, of which 1,647 were TPO compatible (20.376%).

#### TPO-enrichment-based secondary driver gene candidate selection axis

The final driver candidate set was the union of the ΔRV top-300 and a complementary TPO-enrichment axis. Genes were assigned to the TPO-enrichment axis when they had at least five canonical coding-intron events, at least three TPO events and a raw gene-specific empirical Fah control probability ≤0.10. For each gene, the empirical probability was calculated as the plus-one-corrected upper-tail proportion of 1,000 panel-size- and animal-bin-matched artificial Fah panels in which the simulated TPO fraction was at least as high as that observed in 3×fOnc3: P=[1+N(f_Fah_ ≥ f_Onc3_)]/1001,where N denotes the number of the 1,000 matched artificial Fah panels in which the simulated TPO fraction for that gene was at least as high as that observed in Onc3.

Within each artificial SB-Fah panel, every selected integration occurrence was independently assigned an F0, F1 or F2 construct frame by categorical sampling with probabilities of 0.45, 0.30 and 0.25, respectively, corresponding to the 18:12:10 µg plasmid mixture used for delivery. TPO compatibility was then determined from the assigned construct frame, integration orientation and the inferred intron frame of the overlapping Ensembl-canonical transcript. Panels in which the gene had no canonical coding-intron event–gene assignment contributed a simulated TPO fraction of zero. This raw probability was used as a predefined secondary screening measure. It quantified enrichment relative to the gene-specific Fah background rather than the magnitude of TPO dominance itself.

25 genes met the criterion for the TPO-enrichment axis; of these, 10 were already members of the ΔRV top-300 list, and 15 were added exclusively through the TPO-enrichment axis, resulting in a final set of the 315 strongest driver candidates.

### SB transposon copy number assessment by qPCR

The cumulative copy number of the 3×fOnc3 transposon vectors was determined by qPCR using hydrolysis probe-based detection. Genomic DNA (100 ng per reaction) was analyzed in three technical replicates using PerfeCTa qPCR ToughMix (Quantabio) on a Rotor-Gene Q real-time PCR instrument (QIAGEN).

The SB transposon was detected using the previously described IRDR-R primer and probe set targeting the right inverted repeat-direct repeat region of the transposon [27]. The mouse TaqMan Copy Number Reference Assay targeting the telomerase reverse transcriptase gene (Tert; Applied Biosystems/Thermo Fisher Scientific, cat. no. 4458368) was used as the endogenous genomic reference.

Cycling conditions consisted of an initial denaturation at 95°C for 7 min, followed by 40 cycles of 10 s at 95°C and 30 s at 60°C. PCR efficiencies were analyzed with Rotor-Gene Q software (QIAGEN). Relative SB transposon copy number was calculated using the ΔCt method [28], with Tert used for normalization.

### Mouse-to-human ortholog assignment

The mouse driver gene candidate panel Top315, the union of ΔRV top300 and 15 TPO-rescue-only genes, comprised 315 genes without restriction by gene biotype. Mouse-to-human orthology mapping was performed for the 243 protein-coding genes (M243), while 72 non-protein-coding genes were not subjected to ortholog assignment. Protein-coding mouse genes were mapped to human orthologs using the Mouse Genome Informatics (MGI) Homology data table (HOM_MouseHumanSequence) as the primary orthology source. MGI Alliance Homology records were used as an offline fallback where no suitable HOM_MouseHumanSequence record was available. Mouse gene symbols and Ensembl identifiers were resolved using MGI marker and gene model reference files. Human gene symbols were checked and updated against HUGO Gene Nomenclature Committee (HGNC) complete and withdrawn symbol tables. Of the M243 protein-coding genes, 228 had a single recommended human ortholog symbol and constituted the human ortholog panel (H228). Downstream human CGC and COSMIC analyses used only these genes with a single recommended human symbol.

### Cancer Gene Census annotation and enrichment analysis

The H228 symbols were annotated against a local export of the COSMIC Cancer Gene Census (CGC), release v104, including both Tier 1 and Tier 2 genes. The background universe comprised 44,997 HGNC-approved human gene symbols. To preserve the HGNC locus-group composition of the query panel, 10,000 random gene sets were sampled without replacement, each containing 226 symbols classified by HGNC as protein-coding genes and two symbols assigned to other HGNC locus groups. CGC genes were counted in each random set. A one-sided empirical enrichment p value was calculated using the plus-one estimator: p = (1 + number of random gene sets with CGC hits ≥ observed CGC hits) / (1 + number of random iterations). The analysis used a fixed random seed to ensure reproducibility. 18 of the 228 human ortholog symbols were annotated in the CGC (7.89%; 17 Tier 1 and one Tier 2 gene). Length-independent composition-matched random gene sets contained a mean of 8.82 CGC genes (median, 9; 95% empirical interval, 4–15), corresponding to a 2.04-fold enrichment and a one-sided empirical p value of 0.0033.

### TCGA MC3 coding-mutation enrichment analysis of the H228 ortholog panel

Somatic coding mutations were analyzed using the public TCGA Multi-Center Mutation Calling in Multiple Cancers (MC3) v0.2.8 mutation annotation format file. Tumor samples were restricted to the cBioPortal TCGA PanCancer Atlas sequenced case lists, yielding 10,443 pan-cancer samples across 32 studies. Only PASS variants, indicating that they passed the MC3 technical quality and artifact filters, annotated with a protein-coding biotype were retained. Current gene symbols were assigned from MC3 HGNC identifiers using the HGNC complete set.

Variant classes were summarized as follows: nonsynonymous variants comprised missense, nonsense, frameshift, in-frame indel, splice-site, translation-start, and nonstop mutations; strict truncating variants comprised nonsense, frameshift, nonstop, and translation-start mutations; and silent variants were analyzed as a synonymous negative control. Gene-level prevalence was defined as the number of unique sequenced tumor samples carrying at least one qualifying variant in the gene divided by the total number of sequenced samples in the corresponding cohort.

The H228 panel contained 228 single recommended human ortholog symbols assigned to protein-coding mouse driver genes. MANE GRCh38 v1.5 protein length was used as a coding-length proxy. One gene, CCDC162P, lacked a MANE protein length and was retained in descriptive H228 outputs but excluded from length-matched inference, resulting in an H227 length-analyzable panel. Eighteen Cancer Gene Census genes were removed in a second analysis, producing an H209 non-CGCpanel.

For each target panel, 10,000 random gene sets of identical size were sampled. Matching was performed within 100 quantile bins of log2-transformed MANE protein length, with each random set containing exactly the same number of genes per length bin as the target panel. The primary control universe comprised current HGNC protein-coding genes with an available MANE protein length, excluding H228 genes. In analyses of the H209 non-CGC panel, CGC genes were also excluded from the respective control universe.

For each random set, median nonsynonymous, missense, truncating, and synonymous prevalence were calculated. One-sided empirical enrichment P values were calculated using a plus-one correction as p = (1 + number of random values ≥ observed value) / (10,000 + 1)’. P values were adjusted by the Benjamini-Hochberg method within each combination of target panel, cohort, and control universe. The minimum attainable empirical P value was approximately 1.0 x 10^-4^.

### Reactome pathway over-representation analysis

Reactome pathway enrichment of the final H228 human ortholog set was evaluated by gene-level over-representation analysis using a frozen Reactome v97 GMT collection. Reactome disease pathways were excluded, and the analysis was restricted to non-disease pathways containing 10–300 unique genes, yielding 1,485 tested pathways. Enrichment was assessed using one-sided hypergeometric tests with a fixed 20,000-gene background, and P values were adjusted jointly across all 1,485 tests using the Benjamini–Hochberg procedure; BH q < 0.05 was considered significant.

For visualization only, nominally enriched pathways (P < 0.05) with at least three overlapping drivers were grouped when they were hierarchically related or belonged to the same top-level Reactome topic and showed a driver-overlap Jaccard similarity of at least 0.50. The most significant pathway represented each cluster. Two individually significant synaptic pathways belonging to the same redundancy cluster—Neurexins and neuroligins and Protein-protein interactions at synapses— were displayed as separate rows linked by a common cluster annotation. ΔRV values, driver ranks, and TPO rescue status were used exclusively for descriptive plotting and did not enter the enrichment statistics.

